# Riparian ecosystem in the alpine connectome. Terrestrial-aquatic and terrestrial-terrestrial interactions in high elevation lakes

**DOI:** 10.1101/035576

**Authors:** Dragos G. Zaharescu, Antonio Palanca-Soler, Peter S. Hooda, Catalin Tanase, Carmen I. Burghelea, Richard N. Lester

## Abstract

Alpine regions are under increased attention worldwide do their role in storing freshwater of high quality and their high sensitivity to climate change - comparable only to the poles. Riparian ecosystems in such regions, integrating water and nutrient fluxes from aquatic and terrestrial environments, host a disproportionally rich biodiversity, despite experiencing severe climate and nutrient restrictions. With climate change rapidly encroaching in the alpine biome, it is important to fully understand how the lake and its surrounding landscape elements sustain such rich ecosystems, before their functional connectivity could be seriously severed.

A total of 189 glacial origin lakes in the Central Pyrenees were surveyed to test how key elements of lake and terrestrial environments work together at different scales to shape the riparian plant composition. Secondly, we evaluated how these ecotope features drive the formation of riparian communities potentially sensitive to environmental change, and assessed their habitat distribution. At each lake plant taxonomic composition was assessed together with elemental composition of water and sediment and ecosystem-relevant geographical factors.

At macroscale vegetation composition responded to pan-climatic gradients altitude and latitude, which captured, in a narrow geographic area the transition between large European climatic zones. Hydrodynamics was the main catchment-scale factor connecting riparian vegetation with large-scale water fluxes, followed by topography and geomorphology. Lake sediment Mg and Pb, and water Mn and Fe contents reflected local connections with nutrient availability, and water saturation of the substrate.

Community analysis identified four keystone plant communities of large niche breadths, present in a wide range of habitats, from (i) damp environments, (ii) snow bed-silicate bedrock, (iii) wet heath, and (iv) limestone bedrock. With environmental change advancing in the alpine biome, this study provides critical information on fundamental linkages between riparian ecosystem and surrounding landscape elements, which could prove invaluable in assessing future biomic impacts.

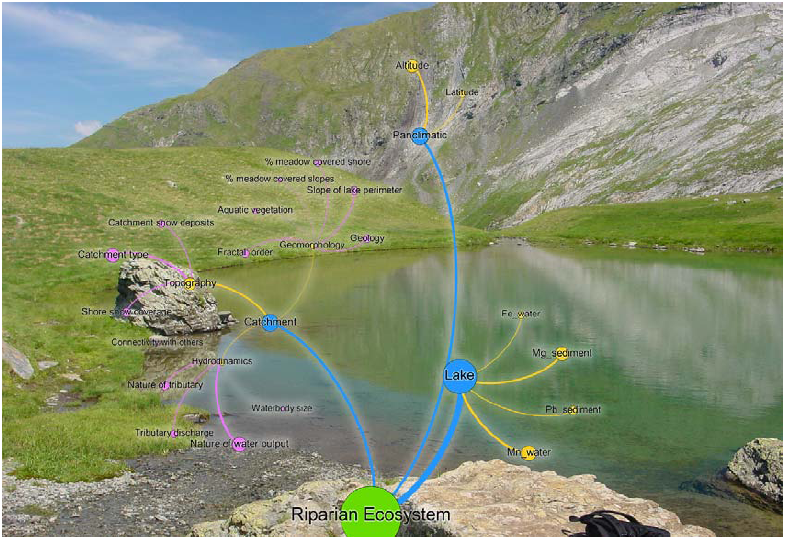
Graphical abstract: Riparian ecosystem of Lake Cardal (0.3ha, 2224m a.s.l) in the Pyrenees National Park (France), with a network diagram of connected landscape elements. Photo by Antonio Palanca-Soler.

## 1. INTRODUCTION

Although they occupy < 24 % of Earth’s land surface, directly and indirectly, mountains provide resources for more than half of the humanity, as well as they contribute > 50 % of total nutrients to the wider biosphere (Price, 2004; Larsen et al. 2014). This is primarily due to an elevated and steep topography, and exposed geology, which create conditions for water precipitation and accumulation, and nutrient release through accelerated bedrock weathering (Larsen and others 2014). Their upper riches - alpine biome, experience the harshest climatic and topographic settings, and a weakly developed soil. These conditions sustain ecosystems strongly connected to the underlying bedrock, which are highly sensitive to small changes in external factors such as atmospheric chemistry (Storkey and others 2015) and climate (Williamson et al., 2009).

Most of the low-lying landforms of the present mountain landscape, including the vast majority of mountain lakes, are the legacy of Pleistocene glaciation (Thornbury, 1969). This produced more than 50,000 remote mountain lakes in Europe alone (Kernan et al., 2009), and > 4000 in the Pyrenees (Castillo-Jurado, 1992). At the interface between terrestrial and aquatic environments, riparian surfaces mediate the fluxes of water, nutrients, and carbon between lakes and surrounding catchments. They also host a disproportionately high diversity of life forms compared to the surrounding landscapes (Gregory et al., 1991, Kernan et al., 2009), and are extensively distributed in the mountain biome (> 797 km only in the Pyrenees; Castillo-Jurado, 1992).

Cross-scale interactions between vegetation, surface properties (including bedrock morphology and geochemistry) and climate determine species distribution in patterns along environmental gradients (Austin and Smith, 1989; Hengeveld, 1990). Baroni-Urbani et al. (1978) introduced the term *chorotype* to define a pool of species with overlapping distributions. Fattorini (2015) revisited the concept and further classified the chorotype into global (for worldwide spatial interactions) and regional. A regional chorotype is assumed to occupy a small geographic area, commonly used as a study area within a biome, and it can be more or less uninterrupted. To group species into chorotypes, cluster analysis is the statistical tool generally preferred (Fattorini 2015). When clustering membership of species cannot be established statistically, they are assumed to follow continuum distributions (Báez et al., 2005).

In the alpine biome, the rough topography and stress severity from low temperature, abrasion by snow and ice, high UV radiation and water-level fluctuations, potentially drive fragmentation of riparian populations into insular communities tightly connected to local resources. Waterbodies isolation could also limit gene flow among such communities. Restrictive influences in plant distribution have been shown over localized areas, due to climate factors such as the type and intensity of precipitation, daily temperature, the frequency of freezing events and their duration (Keller et al., 2005), as well as slope orientation and altitude (Baker, 1989).

Climate change, particularly precipitation, air temperature, freezing line and snow cover can greatly influence the thermodynamics and geochemistry of high altitude catchments (Thompson et al., 2005; Parker et al., 2008; Zaharescu et al. 2016a), and consequently their riparian communities. With species populations of many mountain ecosystems reaching their tipping points (Kreyling et al., 2014; Buma et al. 2016), it becomes critically important to better define the breadth and strength of their natural connection with the supporting physicochemical template (ecotope) at both, local-scale and the broader landscape, before their possible loss due human encroachment and climate change. We use the term connectome (first introduced by Sporns, 2006, and Hagmann, 2005 in neurosciences) to denote the functional linkages between a lake ecosystem and the wider environment, as it offers a natural way to understand ecosystem interactions.

Research exploring the connection between riparian ecosystem and its supporting physicochemical template in the conceptual framework of ecotope is rare. It has largely been conducted at low altitudes, e.g. focusing on local scale alterations in hydrological and habitat disturbance affecting riparian communities (Merritt et al., 2010). Related work in high elevation watersheds has recently quantified the relative contribution of geomorphic characteristics to predict the type of riparian plant communities and species abundances at different scales (Engelhardt et al., 2015). More recently, we conceptualized the ecotope model (Zaharescu et al., 2016b), and explored its influence on the benthic ecosystem (Zaharescu et al., 2016c) in alpine lakes. Here we further assess how major ecotope properties interact at different scales to shape the structure of riparian plant communities in an alpine region of limited human impact.

The motivation behind this study was to address three questions in ecology, key in attaining a mechanistic understanding of how high-elevation ecosystems develops in relationship with their physical template, and their potential response to environmental change. They were: (i) to assess the cross-scale linkages between catchment surface properties, lake, and riparian ecosystem, and evaluate their strength; (ii) to identify keystone plant communities and lakes potentially sensitive to environmental change, and (iii) to evaluate their habitat distribution along environmental gradients. We postulate that because of the extreme geoclimatic setting, riparian ecosystems in alpine regions are more strongly connected to local lake variables than to larger scale gradients. This, in turn, would create local indicator communities highly susceptible to environmental change.

The location of the Pyrenees at the intersection of four large biogeographical regions in Europe (Atlantic, Continental, Mediterranean, and Alpine), makes them richer in biodiversity compared to similar areas such as the Alps, with a high proportion (± 11.8 %) of endemic plant species (Gómez et al., 2003). They are, therefore an excellent candidate for such study.

## 2. METHODS

### 2.1 The area

The Pyrenees are a relatively young mountain chain in SW Europe and a natural barrier between Spain and France. Their topography was carved mostly during the last glaciation 11,000 years ago, which left an abundance of lakes and ponds in cirques and valleys. The lakes, in different stages of ecosystem evolution, are more abundant on the steeper French side, which generally receives more precipitation.

The study area extended from - 0°37’35” to 0°08’19” E and 42°43’25” to 42°49’55” N in the axial region of the Pyrenees National Park, France (Fig. 1). The geology is dominated by granitic batholiths, surrounded by old metamorphic and sedimentary materials, including slates, schist, and limestone. The hydrology is broadly shaped by Atlantic influences, which feed > 400 lakes and ponds in the park. A great number of lakes are drained by temporary torrents and permanent streams, which converge into major valleys, though isolated waterbodies and karstic systems are not rare. The valleys generally follow an S - N direction. Some of the big lakes at lower altitudes were transformed into reservoirs and are used for hydropower and as freshwater reserves of high quality.

**Figure 1.**
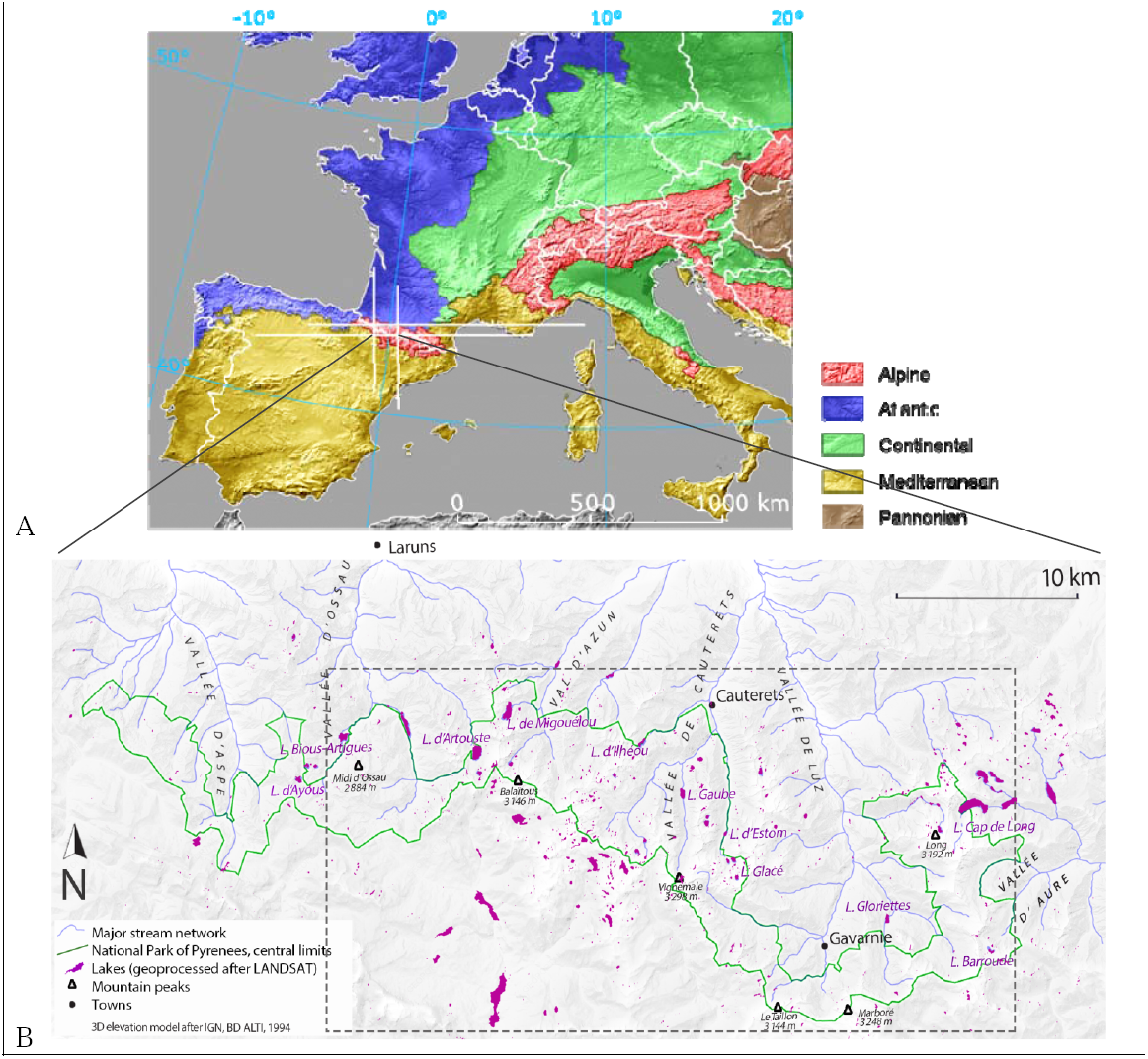
Biogeographical regions of W Europe (A, after EEA, 2001), with the location study area (B) in the axial part of Pyrenees National Park, France. Surveyed lakes, enclosed by a dash-line box, were within the boundaries of the park.

### 2.2 Sampling strategy

A total of 189 high-altitude lakes and ponds, ranging from 1161 to 2747 m a.s.l. was visited in July of 2000, 2001 and 2002. Lake sampling was designed to cover the majority of water bodies in the central region of the park (of highest altitudes) in the shortest possible period, which corresponded to the summer phenology.

Each lake was characterized according to riparian vegetation composition and a range of catchment physical and chemical attributes (Table S1). Ocular identification of species around each waterbody was recorded in the field using Grey-Wilson and Blamey (1979), Fitter et al. (1984) and García-Rollán (1985) keys. Hard to identify plants were collected and transported in a vasculum to the laboratory for identification. They were thereupon identified using Flora Europaea (available online at http://rbg-web2.rbge.org.uk/FE/fe.html).

At each location, hydrological (tributary discharge, nature and size of tributary and output), geomorphological (bedrock geology, % slope of lake perimeter, fractal development level, % shore and near-catchment slope covered by meadow and aquatic vegetation) and topographical (catchment type, catchment and shore snow coverage and connectivity with other lakes) attributes were visually inspected and scored according to dominant units (Table S1). Geolocation, i.e. altitude, latitude, and longitude was recorded at each lake using a portable GPS device.

To test for relationships between lake chemistry and riparian vegetation composition, < 3 cm depth littoral sediments and water ± 5 m off the littoral (for small waterbodies, the distance was less) were sampled using a polyethylene trowel. The sediments comprised fragmented rocks, coarse sands, and fine silts. As the chemical composition of the fine sediment fraction is the most likely to relate to riparian vegetation, i.e. they are either source or sink of the bioavailable element fraction), sampling deliberately targeted this fraction. To assure sample homogeneity, at each lake the sample comprised roughly 5 randomly selected subsamples. All sediment and water samples were kept at < 4°C until laboratory analysis.

Water pH and conductivity were recorded on site, at the surface and bottom of the lake from samples taken with a Teflon bottom water sampler. Portable pH and conductivity probes were used in this case.

### 2.3 Sample preparation for major and trace element analysis

The sediment samples were dried at 40 °C for two days and sieved through a 100 μm sieve. Trace and major element contents were characterized by X-ray fluorescence spectrometry (XRF). A 5 g portion of the sample was prepared as lithium tetraborate melt for the determination of major (Na, Mg, Al, Si, P, S, Cl, K, Ca, Ti, and Fe) and trace (V, Cr, Mn, Co, Ni, Cu, Zn, As, Rb, Sr, Ba and Pb) elements. Results are expressed in % mass-mass and mg kg^-1^, respectively for major and trace elements. Fusions were performed in Pt–Au crucibles. Calibration and quality control analyses were carried out using replicated certified reference materials from National Research Council of Canada, NRCC (SO-3, SO-4, HISS-1, MESS-3 and PACS-2, soils and sediments) and from South Africa Bureau of Standards, SACCRM (SARM 52, stream sediment). Additionally, a given sample was analyzed several times during the analysis run. The analysis was highly reliable, with the recovery figures for the reference materials being within an acceptable range for all major elements (± 10 %). Percent coefficient of variation (% CV) between replicates was < 5 % and % relative standard deviation, RSD (1 σ) between measurements of the same sample < 2 %.

Total C and N contents were simultaneously determined by flash combusting 5 mg dried sediments in a Carlo Erba 1108 elemental analyzer following standard operating procedure (Verardo et al., 1990).

Water samples were prepared for analysis by filtering through 0.45 μm cellulose nitrate membrane followed by acidification to 2 % with ultrapure Merck nitric acid. The acidified samples were analyzed for Li, B, Na, Mg, Al, K, Ca, Mn, Fe, Ni, Cu, Ga, Se, Sr, Ba, Rb and Pb by inductively coupled plasma - optical emission spectrometer (ICP-OES) using standard ICPMS/ OES operating conditions. The results are expressed in mg L^-1^. The analysis, following standard procedures and QA/QC protocols, were performed at the University of Vigo’s Centre for Scientific and Technological Support (CACTI).

### 2.4 Statistical procedures

#### 2.4.1 Principal component analysis to summarize ecotope factors

Principal component analysis (PCA) was used to reduce the multiple catchment-scale variables to a limited number of composite factors (principal components, PCs), which represent major ecotope properties being investigated, according to the procedure developed in Zaharescu et al. 2016b (Table 1). The regression factor scores of the principal components (extracted after an orthogonal *Varimax* rotation) were then used as explanatory variables in the further analysis of vegetation data. This analysis was performed in PASW (currently IBM-SPSS) statistical package for Windows.

#### 2.4.2 (Multidimensional) Fuzzy Set Ordination to quantify riparian drivers

To understand the potential effects of catchment gradients on vegetation species composition we used Fuzzy Set Ordination (FSO) followed by a forward - stepwise multidimensional FSO (MFSO), both run on a distance matrix of species incidence data.

Introduced by Roberts (1986), FSO is a more natural alternative to constrained ordination methods, e.g. CCA and RDA. Compared to the classical theory in linear algebra, where cases are either in or out of a given set (0 or 1), in FSO cases are assigned partial membership (fuzzy) values ranging from 0 to 1 that denote their degree of membership in a set (Roberts, 2008). Likewise, species responses to environmental factors are generally not limited to a certain function; they can be, for example, nonlinear or discontinuous. FSO, therefore, is a generalized technique (Roberts, 1986), which overcomes this problem and includes the types of ordination that ecologists are more familiar with, such as gradient analysis (Whittaker, 1967) and environmental scalars ordination (Loucks, 1962). Thus, in fuzzy logic applications, the results can facilitate the expression of natural rules and processes.

First, a distance matrix of species incidence was calculated. For the binary data (presence-absence) considered herein, we used Sørensen similarity index, as suggested by Boyce and Ellison (2001). This gave a measure of similarity between sites based solely on species composition. This was followed by one-dimensional FSO, taking distance matrices of species as response variables and the environmental variables as explanatory variables. FSO also requires that the environmental variables be as much uncorrelated as possible (Boyce, 2008). A number of landscape variables showed a strong correlation. Their summarized version, i.e. the PC regression factor scores from prior PCA (Table 1), were therefore used as explanatory variables in FSO. By default, the principal components of PCA computed with an orthogonal (e.g. Varimax) rotation are uncorrelated, therefore, suitable for this approach.

**Table 1.** Association of lake catchment variables into three composite factors. Variables are displayed in the order of correlation with the principal components (PC). Highest correlation of a variable with any of the components is in bold. PC1 was interpreted as hydrodynamics; PC2, geo-morphology, and PC3, topography formation. Variable values are in Table S1.

|  | Principal component |  |  |
| --- | --- | --- | --- |
|  | PC 1 | PC 2 | PC 3 |
| Tributary discharge | **0.92** | 0.04 | 0.02 |
| Nature of tributary | **0.90** | 0.02 | 0.01 |
| Nature of water output | **0.87** | -0.17 | 0.07 |
| Waterbody size | **0.52** | -0.38 | 0.05 |
| % meadow covered slopes | -0.07 | **0.72** | -0.37 |
| % meadow covered shore | 0.21 | **0.68** | -0.24 |
| Slope of lake perimeter | 0.30 | **-0.67** | -0.03 |
| Geology | -0.23 | **0.60** | 0.07 |
| Aquatic vegetation | -0.16 | **0.58** | -0.22 |
| Fractal order | 0.07 | **0.50** | 0.08 |
| Catchment snow deposits | 0.09 | -0.10 | **0.86** |
| Catchment type | 0.05 | 0.07 | **0.79** |
| Shore snow coverage | -0.11 | -0.11 | **0.75** |
| Connectivity with others | 0.39 | -0.36 | **0.52** |
| Total Eigenvalue (rotated) | 3.07 | 2.69 | 2.46 |
| % of variance explained | 21.96 | 19.24 | 17.59 |
| Cumulative % | 21.96 | 41.20 | **58.79** |

Rotation method: Varimax with Kaiser normalization.

To remove potential covariance between factors and better quantify the effect size of each factor on riparian vegetation, a multidimensional FSO (MFSO) was run on factors with correlation coefficient with species distance matrix (r) > 0.3 (p < 0.05). This allowed a more accurate multidimensional interpretability of the results. Statistically, MFSO first performs an FSO on the variable accounting for most of the variation. Then, the residuals from that fuzzy ordination are used with the next most important variable, and the process is repeated until no more variables are left. Therefore, unlike the widespread ordination methods used in ecology, e.g. Canonical Correspondence Analysis (CCA) and distance-based redundancy analysis (DB-RDA), in MFSO each variable selected by the model can be considered as an independent axis, and only the fraction of axis membership values which is uncorrelated with previous axes is included into the model (Roberts, 2009a). Moreover, MFSO is expected to perform better than other unconstrained ordination methods on more complex datasets, as it is less sensitive to rare species and noise in environmental factors (Roberts, 2009a) - therefore better reflecting the average community behavior.

The effect magnitude of each environment variable on plant species composition was assessed visually by the relative scatter attributable to that variable, and numerically by its increment in the correlation coefficient. Similar to regression modeling, if an axis is influential in determining the distribution of vegetation, then one should be able to predict the values of that variable based on species composition (Roberts, 2009b). A “step-across” function was used on MFSO results, to improve the ordination with high beta diversity, i.e. for sites with no species in common (Boyce and Ellison, 2001).

The significance of the matrix correlation coefficient between environmental variables and species composition was established by permuting the rows and columns of one of the matrices 1000 times in both, FSO and MFSO, recalculating the correlation coefficient and comparing the observed matrix correlation coefficient with the distribution of values obtained via permutation.

FSO and MFSO were computed with FSO (Roberts, 2007a) and LabDSV (Roberts, 2007b) packages, while the step-across function was computed with VEGAN package (Oksanen et al., 2009), R statistical language and environment.

#### 2.4.3 Network diagram

A conceptual diagram summarizing the linkages between riparian vegetation and surrounding landscape elements, and their magnitude of influence was assembled in the open-source visualization tool Gephi 0.9.1 (https://gephi.org; Bastian et. al. 2009) using Yifan Hu layout. Gephi is a highly interactive visualization platform capable of displaying the relationships between nodes of a semantic network based on object abundances or objects weight. It allows users to intuitively discover patterns, isolate network structures, and singularities, and derive hypotheses in social and biological networks analyses. We used the independent (incremental) *r* values derived by MFSO as variable weights. Likewise, a multiple regression with Akaike Information Criterion model selection (Automatic Linear Modelling option in IBM-SPSS) provided the magnitude of influence each landscape variable has in the composite landscape factors previously summarized by PCA.

#### 2.4.4 Indicator community analysis

The riparian vegetation composition (incidence data) was analyzed for species association into chorotypes, i.e. species with significant co-occurrence patterns. First, the lakes were grouped on the basis of shared species. For this, a clustering procedure (Pair-Group Method using the Arithmetic Averages (PGMA) using flexible linkage (beta) parameter, with beta = -0.2) was computed using the Sørensen distance matrix of species incidence. This allowed selecting the most appropriate clustering for dendrogram nodes cut.

The selected clusters were subsequently assigned code numbers into a new categorical variable. This variable was used as a grouping variable in Indicator Species Analysis (Dufrene and Legendre, 1997), to determine plant species with significant affinity to the lake categories, i.e. species of similar ecological preferences. An indicator community comprises species that are most characteristic in the riparian zone of lakes of that type. The higher the indicator value is, the greater is the species affinity to a lake type.

Furthermore, the selectivity of the resulting vegetation communities to different ecotope features was tested by box-plotting them against environmental gradients. Sørensen similarity matrix was computed with ADE4 (*dist.binary* function; Thioulouse et al., 1997), cluster and boxplot analyses with CLUSTER (*agnes* and *boxplot* functions, respectively; Kaufman and Rousseeuw, 1990), Discriminant Analysis with FPC (*plotcluster* function; Hennig, 2005) and Indicator Species Analysis with LabDSV (*indval* function; Dufrene and Legendre, 1997) packages for R statistical language (R Core Development Team, 2005); available online at http://cran.rproject.org/.

## 3. RESULTS AND DISCUSSION

### 3.1 Major catchment-scale variates

Interactions between the lake, catchment, and larger geographical gradients create major driving forces which shape the development of land-water interface ecosystems. PCA extracted three major components, which accounted for more than 58 % of the total variance in lake catchment characteristics (Table1).

The first principal component (PC1) was interpreted as hydrodynamics and was formed by tributary nature and discharge, type of water output and waterbody size (Table1). The second component (PC2) characterizes the major bedrock geo-morphology, i.e. geology, shore sloping, % of slope and shore covered by meadow, fractal order (riparian development) and the presence of aquatic vegetation. The third PC represents topography, i.e. catchment type, visible connectivity with other lakes, and catchment and shore snow deposits. The three composite factors were, therefore regarded as major landscape drivers expected to influence the lake riparian ecosystem development.

### 3.2 Influence of lake, catchment, and pan-climatic drivers on riparian vegetation

A single-dimensional FSO, used as the initial step in evaluating the cross-scale linkages between catchment surface properties, lake, and vegetation composition (first objective), clearly identified altitude to exert the largest influence, followed by water Mn and Fe concentration, sediment Mg contents, horizontal gradients latitude and longitude, and catchment-scale variables topography formation and hydrodynamics (Table 2). The effect size of each of the factors correlating with vegetation composition at r > 0.3 (p < 0.05; Table 2) were quantified by MFSO to remove potential covariation. The results are displayed in Table 3 and are discussed in the following sections.

**Table 2.** One-dimensional fuzzy relationships between riparian vegetation species composition and environmental factors in the Pyrenees lakes. Factor superscripts: (a) geoposition, (b) landscape (Table 1), (c) sediment chemistry, and (d) water chemistry. Correlations between factors and apparent factors predicted by vegetation are listed in descending order. Factors with correlations > 0.3 (in bold) were retained for further MFSO analysis. *P* represents the probability after 1000 permutations.

| Variable | *r* (Pearson) | *P* | Variable (*continued*) | *r* (Pearson) | *P* |
| --- | --- | --- | --- | --- | --- |
| <sup>a</sup>Altitude | **0.855** | 0.001 | <sup>c</sup>Sr | -0.005 | 0.545 |
| <sup>a</sup>Latitude | **0.695** | 0.001 | <sup>c</sup>Na | -0.020 | 0.439 |
| <sup>a</sup>Longitude | **0.636** | 0.001 | <sup>c</sup>Ti | -0.107 | 0.540 |
| <sup>b</sup>Topography (PC3) | **0.644** | 0.001 | <sup>c</sup>Rb | -0.164 | 0.624 |
| <sup>b</sup>Geo-morphology (PC2) | **0.603** | 0.001 | <sup>c</sup>Al | -0.443 | 0.900 |
| <sup>b</sup>Hydrodynamics (PC1) | **0.442** | 0.001 | <sup>d</sup>Mn | **0.751** | 0.001 |
| <sup>c</sup>Mg | **0.712** | 0.001 | <sup>d</sup>Fe | **0.730** | 0.001 |
| <sup>c</sup>Pb | **0.515** | 0.003 | <sup>d</sup>Conductivity (surface) | **0.584** | 0.001 |
| <sup>c</sup>Ca | **0.510** | 0.004 | <sup>d</sup>Conductivity (bottom) | 0.545 | 0.001 |
| <sup>c</sup>Cu | **0.501** | 0.007 | <sup>d</sup>Al | **0.531** | 0.014 |
| <sup>c</sup>Co | **0.497** | 0.006 | <sup>d</sup>Cu | **0.465** | 0.009 |
| <sup>c</sup>Ba | **0.484** | 0.007 | <sup>d</sup>pH (bottom) | **0.307** | 0.002 |
| <sup>c</sup>Ni | **0.432** | 0.018 | <sup>d</sup>pH(surface) | 0.257 | 0.002 |
| <sup>c</sup>Mn | **0.405** | 0.024 | <sup>d</sup>K | 0.254 | 0.108 |
| <sup>c</sup>Fe | **0.362** | 0.037 | <sup>d</sup>Na | 0.204 | 0.170 |
| <sup>c</sup>Zn | **0.361** | 0.033 | <sup>d</sup>B | 0.177 | 0.089 |
| <sup>c</sup>C | **0.351** | 0.032 | <sup>d</sup>Pb | 0.130 | 0.272 |
| <sup>c</sup>Si | **0.337** | 0.046 | <sup>d</sup>Ba | 0.108 | 0.248 |
| <sup>c</sup>N | **0.324** | 0.036 | <sup>d</sup>Sr | 0.088 | 0.293 |
| <sup>c</sup>Cr | **0.309** | 0.069 | <sup>d</sup>Se | 0.057 | 0.340 |
| <sup>c</sup>V | 0.210 | 0.130 | <sup>d</sup>Ni | -0.010 | 0.482 |
| <sup>c</sup>C/N | 0.114 | 0.145 | <sup>d</sup>Ga | -0.020 | 0.445 |
| <sup>c</sup>S | 0.112 | 0.249 | <sup>d</sup>Li | -0.030 | 0.462 |
| <sup>c</sup>As | 0.110 | 0.298 | <sup>d</sup>Mg | -0.101 | 0.590 |
| <sup>c</sup>K | 0.029 | 0.342 | <sup>d</sup>Ca | -0.234 | 0.746 |
| <sup>c</sup>P | 0.013 | 0.394 | <sup>d</sup>Rb | -0.350 | 0.841 |
| <sup>c</sup>Cl | -0.001 | 0.418 |  |  |  |

#### 3.2.1 Large vertical and horizontal gradients

Table 3 and Fig. 2a show the environmental factors in order of their independent correlation with the apparent factors predicted by vegetation composition. A two-axes solution resulted from MFSO, with altitude and latitude reliably predicting riparian plant composition (cumulative *r* = 0.65). Altitude, the most influential, is a classical vertical driver of ecosystem development along the alpine climate gradient. The latitudinal variability in the study area (the secondly most important in the model) is relatively narrow. However, its location under four major biogeographical influences, namely Atlantic and Continental from the N and NW, Mediterranean from SE, and the local alpine gradient (Fig. 1) implies that the study area was sufficiently large to capture in its riparian ecosystem macro-regional transitions. Longitude, although showed an individual relationship with vegetation (Table 2), its effect seemed to be a covariate in the multivariate model (Table 3).

**Table 3.**
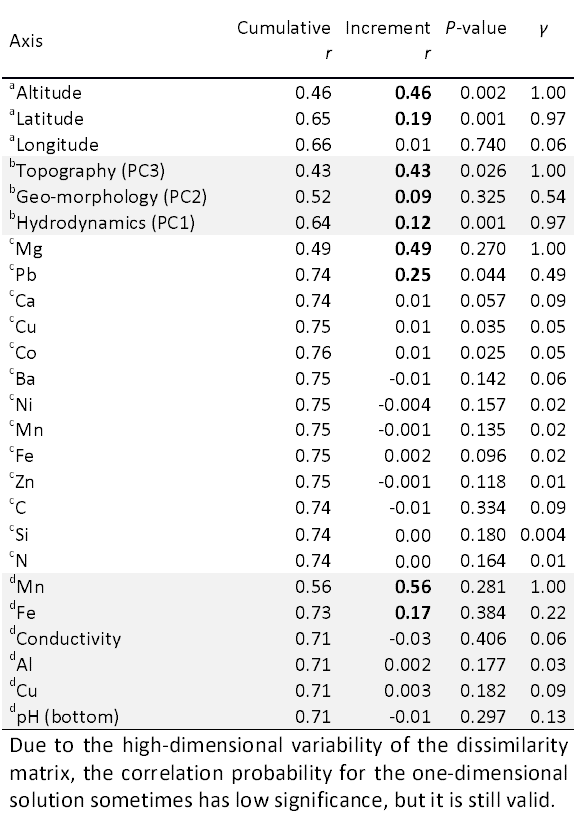
Independent effect of ^a^ geoposition, ^b^ catchment, ^c^ sediment and ^d^ water chemistry factor on riparian vegetation composition, as given by MFSO. Figures for geoposition, catchment and water characteristics resulted from MFSO with step-across function. *γ* (gamma) = a vector of the independent variance fraction of a factor (MFSO axis). Factors with the highest independent influence in the model, in bold, are listed in order of their weight in the model.

**Figure 2.**
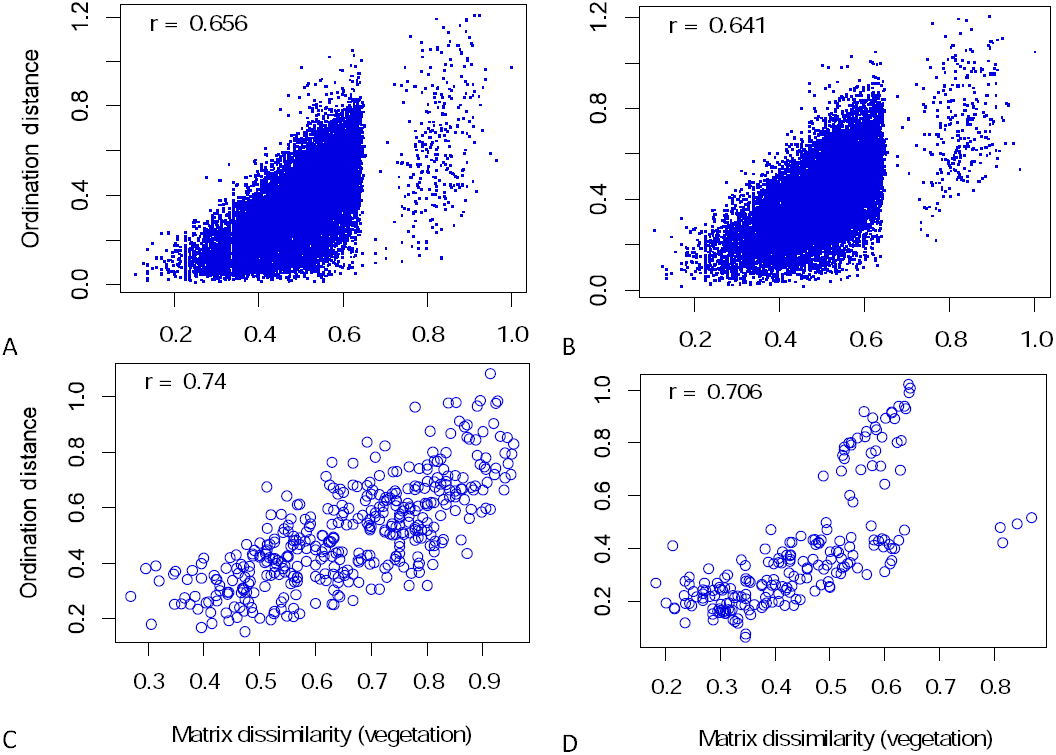
Multidimensional Fuzzy Set Ordination (FSO) depicting the effect of **(A)** large geographical gradients altitude, and latitude, **(B)** catchment factors topography (PC3), hydrodynamics (PC1), and geo-morphology (PC2), **(C)** sediment chemistry, and **(D)** water chemistry, on riparian plant composition. MFSO in A and B were improved by using a step-across function. Variables were introduced in the model in the order of decreasing Pearson fuzzy correlation (*r*; Table 3) with plant dissimilarity matrix. Number of permutations = 1000.

#### 3.2.2 Catchment geomorphological and hydrological elements

MFSO of catchment-scale variables produced a three-dimensional solution, with topography formation largely dominating hydrodynamics and geo-morphology in its influence on vegetation structure (cumulative *r* = 0.64; Table 3 and Fig. 2b). Since these factors are composite (Table 1), it means that catchment type - a legacy of the last glaciation, with local effects from snow cover and lake connectivity (important in propagule dispersion) create variable microclimates, which allow for colonization by different species. For instance, snow at valley head would last longer around lakes and create longer wet conditions than on mountain slopes or passes, which would experience earlier sun exposure. Previous research in alpine regions corroborates these findings and shown that topography and its effect on snow coverage can control terrestrial vegetation (Keller et al., 2005), a consequence of topography interaction with climate variables radiation, precipitation, and wind (Körner, 1999).

Hydrodynamics added to the influence of previous axis (Table 3 and Fig. 2b), through the nature and discharge of tributaries, and associated effects from waterbody size and nature of output (Table 1). These variables control nutrient and sediment fluxes from the drainage basin, and nutrient transfer from lake area to the riparian zone. It implies that stream - fed medium to large lakes (Table 1) host significantly different vegetation than the shallower precipitation - fed ponds.

Although its separate influence was high (Table 2), bedrock geology (with nested effects of slope, vegetation coverage and shore development, PC2), represented the smallest independent driver of riparian vegetation, most likely due to its covariation with topography (Table 3 and Fig. 2b). Geology is known to influence the establishment of vegetation through its control on substrate chemistry and niche formation (Kovalchik and Chitwood, 1990). The bedrock of the area presents an igneous core (granitic) in its central part, flanked by sedimentary and metasedimentary materials. Granitic geomorphology, which is more resistant to weathering, is associated with steep slopes, low vegetation coverage, less developed riparian zones (lower fractal order; Table 1), and contributes fewer nutrients to the lake. Conversely, more reactive bedrock such as limestone sustains a more chemically rich environment, better developed riparian zones (higher fractal order), a more stable terrain with more vegetation coverage (Table 1). Together with topographic and hydrologic particularities, the two geological substrates sustain different riparian species.

#### 3.2.3 Sediment chemistry. Indicator elements

At lake scale, results of the MFSO of riparian vegetation composition and sediment major and trace element contents resulted in a bi-dimensional solution, with Mg and Pb reliably predicting species composition (cumulative *r* = 0.74; Table 3 and Fig. 2c). Catchment lithology, namely the geological structure, the proportion of rock types, their mineralogy, chemistry, and weathering resistance, is tightly connected to the chemistry (major and trace elements and organic matter) of high altitude water bodies through cross-ecosystem fluxes of sediment and water (Lewin and Macklin, 1987; Zaharescu et al, 2009). Magnesium, a component of the photosynthesis molecule chlorophyll, and in many growth enzymes, is an essential macronutrient in green plants. Soils developed on alkaline bedrock (limestone) generally contain more Mg (≈ 0.3 - 2.9 %) than those on more acidic substrates (granite or sandstone; ≈ 0.01 - 0.3 %; Beeson, 1959). Our results further show that plant response to bedrock-derived Mg can determine their community composition in nutrient-poor high elevation environments.

The influence of Pb is rather surprising. One possible explanation is that it reflects the distribution of plants using mycorrhizae (legumes) since mycorrhizal symbiosis has been related with increased Pb uptake under low soil metal concentrations (Wong et al., 2007). Or, plant’s natural sensitivity to Pb (Kabata-Pendias and Pendias, 2001) could have also changed their composition along a natural Pb - stress gradient. It is known that in metamorphic areas of the central Pyrenees, Pb is an abundant mineral constituent (Catalan et al., 2006; Zaharescu et al., 2009).

The low independent effect of other essential elements is likely due to their mineral cooccurrence with Mg and Pb (Spearman correlation coefficient of Mg with Ca, Fe, Cr, Mn, Co, Ni, Cu, Zn, As and Ba r > 0.38, and of Pb with Cu and Zn r > 0.47; P < 0.05). Nonetheless, Mg and Pb merit further mechanistic examination into why they are the dominant indicators of riparian vegetation.

#### 3.2.4 Water chemistry. Indicator elements

The MFSO of riparian vegetation composition and selected water variables, identified Mn and Fe as major influential axes (cumulative *r* = 0.73; Table 3 and Fig. 2d). Besides collecting weathering solutes, water in snow packs affects plant growing season, as well as it controls the temperature and the amount of oxygen reaching the ecosystem. Iron and Mn are major redox players in soil and sediment, and varying water table can modify their solubility and uptake by plants (Alam, 1999). For instance, in the water-saturated soil, biotic respiration drives reduction processes, which affects plant performance not only by preventing nutrient (Mg, Ca and Fe) uptake but also by restricting root development (Couto et al., 1983). Differential response of vegetation to the build-up of soluble Fe and Mn have been suggested in species ecology and habitat distribution (Alam, 1999), and it has been reported for grasses, legumes (Couto et al., 1983) and trees (Good and Patrick, 1987).

Variable moisture and flooding of the riparian ecosystem by lake water are common in alpine lakes, being regulated by snow thaw, and frequency and volume of summer storms. The plant compositional change along Mn and Fe gradients herein are indicative of such process (a nested effect of topography; Table 1), with higher moisture near lakes at valley head and drier conditions on slopes and mountain passes. Unsurprisingly, water pH, conductivity and a number of elements appeared to co-vary with Fe and Mn in the multivariate solution (Tables 2 and 3).

**Figure 3.**
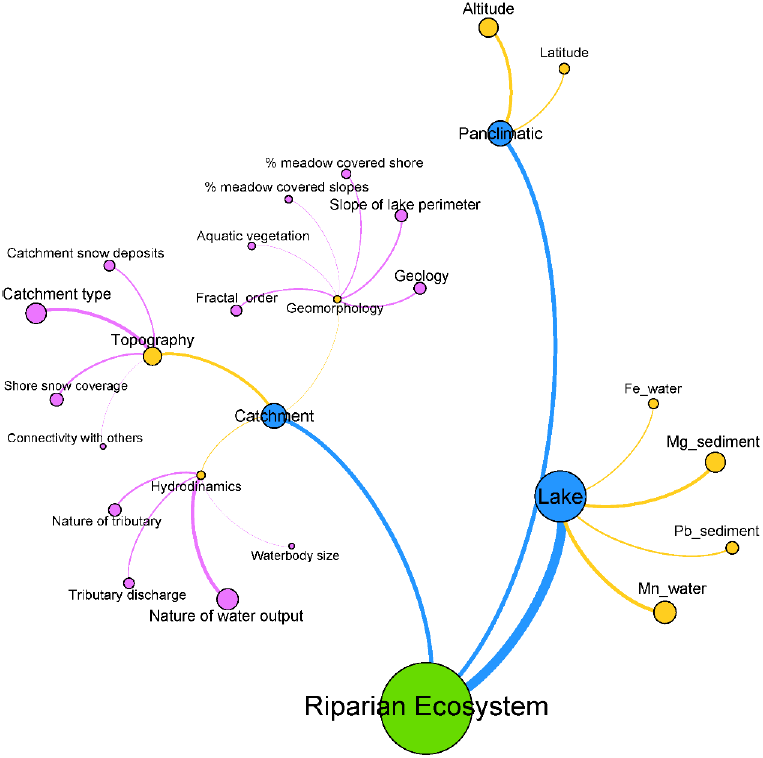
Conceptual network diagram showing the linkages between riparian ecosystem of a typical alpine lake and ecotopic features of the lake, catchment, and large pan-climatic gradients. Nodes and label sizes, as well as connections thickness, are proportional to the magnitude of their influence on target nodes. Connections borrow the colors of source nodes and represent different layers of organization in the model.

#### 3.2.5 Conceptual network diagram

A conceptual network diagram illustrating the connection between riparian ecosystem and ecotopic features at the lake, catchment, and pan-climatic scales is presented in Fig. 3. Important linkages are established between species presence and lake sediment chemistry and water redox condition (reflecting catchment flows of water and nutrients), as well as with the nature of lake source and draining, shoreline development, catchment type and geology, and the large-scale altitudinal and latitudinal gradients. These landscape elements, operating at different scales in their influence on riparian species, are to be sought in future environmental change studies.

**Figure 4.**
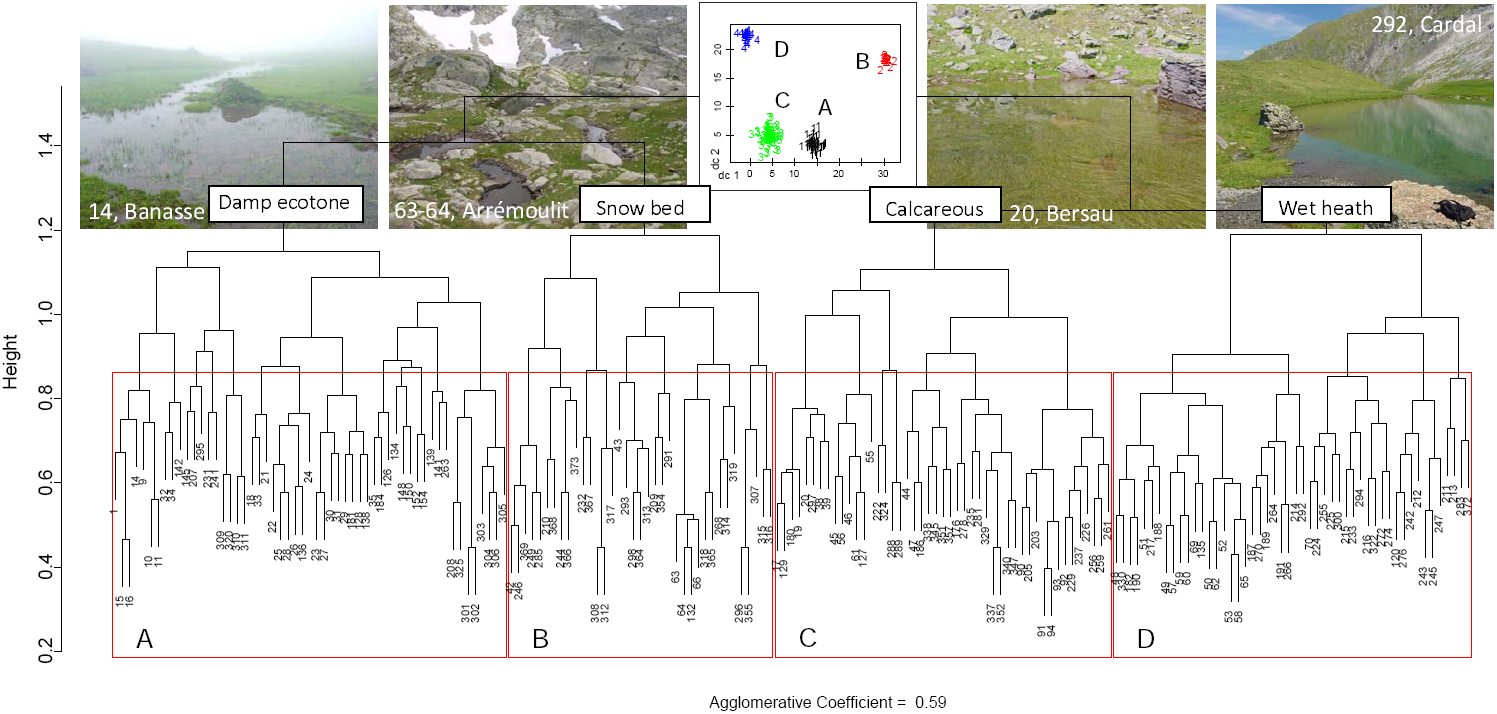
Dendrogram showing lake ecosystem types (clusters) based on similarity in riparian vegetation composition, together with representative examples. A plot of the clusters in discriminating space (inset) shows an effective clustering. Plant communities indicative of lake/ ecosystem types are in Table 4. N = 189 lakes and 166 plant species. Plants are listed in List S1, while lakes in Zaharescu et al., 2016b (supplementary Information).

### 3.3 Indicator communities and their environmental preference

#### 3.3.1 Community analysis

To test whether the restrictive high altitude environment determined the formation of keystone plant communities (i.e. chorotypes; second objective), PGMA cluster analysis was used on species presence data. The results revealed a good grouping (agglomerative coefficient = 59 %), with the 189 lakes classified into four types (Fig. 4). Of the 168 total plant species (List S2), 79 formed four chorotypes, present in the four lake environments. Table 4 shows the species with significant co-preference for each lake type and their probability of group membership. Plant communities A, B, and D yielded the highest degree of confidence (Fig. 4).

Type A lakes (Fig. 4 and Table 4) contains mostly species of damp ecotones, such as bog-associated species with *
*Sparganium, Ranunculus, Chara, Sphagnum* moss, Selaginella* fern, sedges (*Carex*) and rushes (*Juncus*) (Table 4). Associated with these are a limited number of plants of drier/stony habitats, including cosmopolites (*Bellis*), nitrogen-fixing legumes (*Trifolium*) and endemics (*Merendera pyrenaica*). The association tolerates a wide range of bedrock chemistry, including acidic (*Sphagnum*), neutral (*Trifoliums*) and basic (*Polygala*) substrates. This dominant community seems, therefore, to reflect an uneven physical template, common on the lake shores of Central Pyrenees.

The second association (type B), smaller than type A, comprises a high proportion of species with affinity to snow bed and a short growing season (*Saxifraga, Veronica, Sibbaldia*), herbaceous shrubs (*Salix*), and ferns (*Cryptogramma*). Most of them are silicophilous, tolerate a low nutrient substrate of different textures, including scree/rocky, grassland, and damp soil. Endemic grass Festuca eskia plotted with the same group.

**Table 4:** Riparian plant groups and their fidelity to lake ecosystems (Fig. 4), as given by Indicator Species Analysis. A species was classified into a group for which its group indicator value was highest and significant. Cluster C had lower significance.

| Cluster A, damp ecotone, $p < 0.05$ | | Cluster D, calcareous, $p < 0.05$ | |
| --- | --- | --- | --- |
| Species | Indicator value | Species | Indicator value |
| *Potentilla erecta* | 0.47 | *Pinguicula vulgaris* | 0.42 |
| *Caltha palustris* | 0.42 | *Gentiana acaulis* | 0.40 |
| *Parnassia palustris* | 0.36 | *Rhododendron ferrugineum* | 0.35 |
| *Thymus serpyllum* | 0.25 | *Primula integrifolia* | 0.30 |
| *Trifolium repens* | 0.23 | *Vaccinium uliginosum* | 0.30 |
| *Hieracium pilosella* | 0.20 | *Trichophorum cespitosum* | 0.29 |
| *Campanula rotundifolia* | 0.19 | *Calluna vulgaris* | 0.26 |
| *Sphagnum* sp. | 0.19 | *Silene acaulis* | 0.26 |
| *Bellis perennis* | 0.18 | *Trifolium alpinum* | 0.25 |
| *Alchemilla vulgaris* s.l. | 0.17 | *Homogyne alpina* | 0.25 |
| *Sparganium angustifolium* | 0.15 | *Soldanella alpina* | 0.22 |
| *Carex echinata* | 0.14 | *Geum montanum* | 0.21 |
| *Juncus filiformis* | 0.13 | *Vaccinium myrtillus* | 0.21 |
| *Anthoxanthum odoratum* | 0.13 | *Hutchinsia alpina* | 0.20 |
| *Carex nigra* | 0.13 | *Armeria maritima alpina* | 0.19 |
| *Cardamine raphanifolia* | 0.11 | *Phyteuma orbiculare* | 0.18 |
| *Merendera pyrenaica* | 0.11 | *Bartsia alpina* | 0.17 |
| *Prunella vulgaris* | 0.11 | *Viola palustris* | 0.17 |
| *Juncus articulatus* | 0.11 | *Geranium cinereum* | 0.14 |
| *Leontodon autumnalis* | 0.10 | *Luzula alpinopilosa* | 0.12 |
| *Ranunculus aquatilis* | 0.10 | *Lotus alpinus* | 0.11 |
| *Selaginella selaginoides* | 0.09 | *Pedicularis mixta* etc | 0.10 |
| *Polygala alpina* | 0.09 | *Thalictrum alpinum* | 0.10 |
| *Carex flava* | 0.09 | *Saxifraga aizoides* | 0.09 |
| *Polygonum viviparum* | 0.08 | *Gentiana lutea* | 0.06 |
| *Carum carvi* | 0.07 |  |  |
| *Galium verum* | 0.07 |  |  |
| *Luzula desvauxii* | 0.07 |  |  |
| *Ranunculus reptans* | 0.07 |  |  |
| *Sanguisorba officinalis* | 0.07 |  |  |
| *Deschampsia cespitosa* | 0.06 |  |  |
| *Chara foetida* | 0.05 |  |  |
| Cluster B, snow bed, $p < 0.05$ | | Cluster C, wet heath, $p < 0.25$ | |
| Species | Indicator value | Species | Indicator value |
| *Gnaphalium supinum* | 0.51 | *Rumex crispus* | 0.04 |
| *Cryptogramma crispa* | 0.47 | *Carex flacca* | 0.03 |
| *Leucanthemopsis alpina* | 0.34 | *Cochlearia officinalis* | 0.03 |
| *Epilobium alsinifolium* etc | 0.28 | *Leontopodium alpinum* | 0.03 |
| *Sibbaldia procumbens* | 0.25 | *Oxytropis campestris* | 0.03 |
| *Kobresia myosuroides* | 0.23 | *Veronica officinalis* | 0.03 |
| *Veronica alpina* | 0.22 | *Callitriche palustris* | 0.02 |
| *Jasione montana* | 0.21 |  |  |
| *Galium pyrenaicum* | 0.19 |  |  |
| *Poa annua* etc | 0.17 |  |  |
| *Doronicum austriacum* | 0.16 |  |  |
| *Saxifraga stellaris* | 0.14 |  |  |
| *Festuca eskia* | 0.12 |  |  |
| *Meum athamanticum* | 0.10 |  |  |
| *Salix herbacea* | 0.10 |  |  |

N = 166 riparian plant species from 189 water bodies.

Riparian community D is a large community comprising wet heath species of Ericaceae shrubs (*Vaccinium* and *Calluna*), accompanied by snow bed plants (*Primula, Soldanella*, and *Bartsia*), sedges (*Trichophorum*), rushes (*Luzula*), and species of wet substrate (Pinguicula and Homogyne). In small number are species of drier habitat (*Gentiana, Hutchinsia,* and *Phyteuma*), and legumes (*Trifolium*). This community tolerates both, siliceous and calcareous substrates.

Community C was the smallest and least common (p < 0.25; Table 4 and Fig. 4). Its members prefer moist-to-dry calcareous banks. Since the rest of the species had no group association, they can be assumed to follow continuum or gradient distributions (Báez et al., 2005).

The identified communities incorporate species from major terrestrial groups (Gruber 1992; Grey-Wilson and Blamey, 1995; Minot et. al, 2007), but with a preference for the wet riparian environment. They are eurytopic associations of large niche breadths, present in a wide range of habitats. This characteristic is what must have allowed them to colonize the harsh and diverse environment surrounding high elevation lakes. The ecological importance of these communities, however, is that they reflect a natural ecosystem condition. Their further study in connection with climate change models is, therefore, imperative to understand how these communities, and the underlying ecotopes, cope with a changing environment.

#### 3.3.2 Environmental preference of indicator communities

To better define the distribution of identified associations along environmental gradients, and understand their habitats (third objective), they were plotted against geoposition and composite catchment variables (Fig. 5).

As shown by the variability in group medians in Fig. 5, plant communities seemed to respond well to both, large horizontal and vertical gradients, and catchment-scale variables. Community A inhabited water - saturated areas surrounding the larger lakes of large fractal (riparian) order and little summer snow cover. They were on the floors and slopes of (meta)sedimentary glacial valleys, at comparatively low altitudes (median ≈ 2100 m a.s.l.). Community B, with a high proportion of snow bed species, grew in elevated granitic topography (e.g. glacial valleys head and mountain pass, ≈ 2400 m a.s.l.), of steep slopes and low fractal development. These habitats had persistent summer snow cover and lower water turnover (rain-fed lakes). Association C established around small lakes of low input/output, in a wide range of altitude and topography (from valley floors to valley heads). Community D, of the narrowest altitude span, was mostly found inhabiting large catchment head lakes with persistent summer snow and high water turnover.

**Figure 5.**
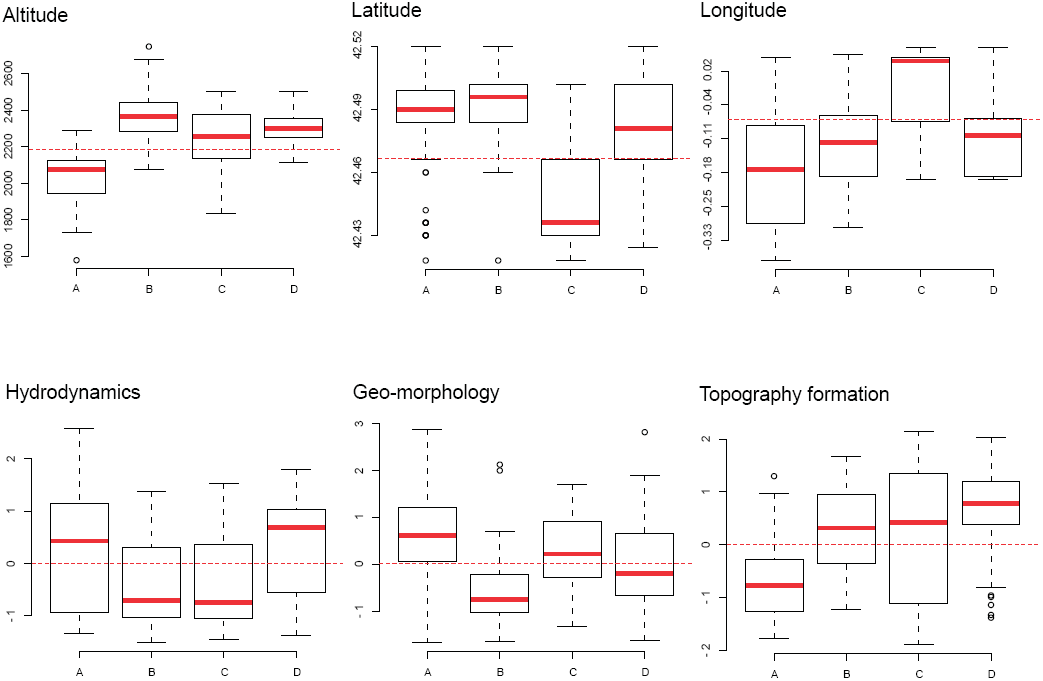
Boxplots showing the distribution of indicator plant communities along geoposition (altitude [m a.s.l.]; latitude [degrees N]; and longitude [degrees E/W, +/-]) and composite catchment gradients. Hydrodynamics factor ranges from small water bodies (-) to large lakes (+), and their associated variables (Table 1); geo-morphology, from igneous and metamorphic (-), to sedimentary (+); topography formation from valley floor and flat topography (-), to valley head (+). Boxplots represent median (red), first (25 %) and third (75 %) quartiles, with whiskers extending to the 5th and 95th percentiles. Horizontal dashed red line is set at plot average.

These results support findings from community analysis and clearly show that none of the evaluated ecotope factors was the sole driver of community establishment in the intricate topography. Rather, a complex pool of microclimatic and geomorphologic conditions worked together to sustain the riparian communities. Since these communities reflect environmental gradients sustaining their formation, their long-term study is necessary, as climate change can affect their distribution through effects on the underlying physicochemical drivers, including precipitation, freezing line, hydrology and weathering fluxes (Zaharescu et al., 2016a).

## 4. CONCLUSIONS

Results showed that alpine lake riparian ecosystems are connected to the lake and terrestrial landscape elements at a variety of scales. Topographical formation left behind by the last glacial retreat, with contemporary effects of snow cover and lake connectivity, are the dominant catchment-scale elements controlling ecosystem development and diversity. Hydrodynamics, with nested contribution from lake size and nature, and size of input and output, was the second most important. The two factors together support the idea that alpine lake ecosystems are not isolated islands in the landscape, but rather interconnected biodiversity nodes controlled by catchment’s physical elements. Geo-morphology, associating geology, shore slope, fractal complexity and vegetation cover greatly covaried with the first two factors, and reflected major geomorphological units in the central Pyrenees, extending from igneous, to metasedimentary and sedimentary materials.

Vegetation relationship with sediment (Mg and Pb) and water (Mn and Fe) chemistry indicated major linkages with nutrient availability and moisture fluctuations in the riparian environment. Superimposed on catchment and local drivers, the ecosystem composition also captured the transition between major biogeographic regions of Europe (effect of latitude and altitude) in an otherwise narrow study area.

The alpine riparian ecotone, connecting complex topography, geology and water regimes, assembled species from both wet and dry environments, which withstand regular flooding. Community analysis identified four such eurytopic groups, i.e. damp ecotone, snow bed-silicates, calcareous and wet heath, of broad niche breadth, characterizing four ecosystem types. These communities preferred different habitat conditions and were distributed along a range of horizontal and vertical gradients in climate, physical and chemical factors.

The strong terrestrial-aquatic and terrestrial-terrestrial connections of the riparian ecosystem at different scales of organisation clearly show how physical elements of the terrestrial surface aggregate into major ecotope drivers, which can predict ecosystem formation, as advanced in Zaharescu et al. (2016b). It remains to be seen how human-induced environmental change affects the functional connectivity between these water-sensitive ecosystems and their supporting ecotope, through its manifold of scales.

## ACKNOWLEDGEMENTS

This study was financially supported by the Pyrenees National Park (France) through a survey grant awarded in 2000-2002. We thank Jorge Milos, Javier Fernandez-Fañanas, Cristina Castan-Lanaspa, David Rodríguez-Vieites, Manuel Domínguez-Rey, Ana Quintillán-Cortiñas, Belén Cirujano-Díaz, Roberto García-Carrera, Jorge Diez-Dieguez, Jorge Rodriguez-Vila, Nicolas Palanca-Castán, Claudia Toda-Castán, Jesús Giraldez-Moreira, Juan Fernández-Rodríguez, Andreea Vasiloiu, Carles Roselló-Vila, Carlos Tur-Lahiguera, Nuria Marti, Maria José Ferrus-Leiva, Julio Palanca-Castán, Bruce Dudley and José Martín-Gallardo for their field and laboratory contribution. We also thank Dave Roberts (Montana State University, USA) and Lasse Ruokolainen (Helsinki University) for their thoughtful statistical suggestions.

## AUTHOR CONTRIBUTION

Design of field research, A. Palanca-Soler; data collection, R.N. Lester, C. Tanase, A. Palanca-Soler and D.G. Zaharescu; study design and manuscript preparation D.G. Zaharescu, P.S. Hooda and C.I. Burghelea. The authors assert no competing interests.

## Supplementary Information

**Table S1.** Landscape and physicochemical variables (and their measured units) estimated for the 189 lakes and ponds in the central region of Pyrenees National Park (France).

| Variable type |  |  |
| --- | --- | --- |
| Numerical | Categorical | Nominal |
| **Latitude**<br>Geographic degrees | **Slope of lake perimeter**<br>Plain, plain in alternation with medium slopes, medium slopes, steep in alternation with medium/plain, steep in >50% of perimeter | **Catchment type**<br>Plain, U shape valley, slope, mountain pass, V shape valley, head of glacial valley |
| **Longitude**<br>Geographic degrees |  |  |
| **Altitude**<br>Meters | **Shore snow coverage**<br>Absent, <10%, 11-50%, >51%, into the water | **Main geology**<br>Conglomerate-sandstone-claystone, limestone (+sandstone-marlstone-schist enclaves), schist (+andesite-sandstone-claystone and granite-limestone), granite (+schist) |
| **Riparian vegetation**<br>Species presence | **Snow deposits in the catchment**<br>Absent, very scarce, scarce, abundant, very abundant |  |
| **Lake sediment C, N, Na, Mg, Al, Si, P, S, Cl, K, Ca, Ti, and Fe**<br>% mass-mass | **Size**<br>Pool (<315±333m<sup>2</sup>), pond (1566±1985m<sup>2</sup>), small lake (9157±10267m<sup>2</sup>), medium size lake (41127±31820m<sup>2</sup>), large lake (91441±37307m<sup>2</sup>) | **Visible connectivity with other**<br>Absent, surrounded by another lake, with another one, in chain |
| **Lake sediment V, Cr, Mn, Co, Ni, Cu, Zn, As, Rb, Sr, Ba and Pb**<br>mg kg<sup>-1</sup> | **Fractal development order**<br>1, 2, 3, 4 | **Nature of tributary**<br>Meteoric, spring, stream/waterfall |
| **Lake water pH (surface and bottom)** | **Tributary discharge**<br>Absent, low discharge, medium discharge, high discharge | **Nature of water output**<br>Absent, temporary, surface-small, surface-medium, surface-large, subterranean, dam output |
| **Lake water conductivity (surface and bottom)**<br>µS cm<sup>-1</sup> | **Aquatic vegetation**<br>Absent, absent but water flooding the grassland, scarce, abundant |  |
| **Lake water Li, B, Na, Mg, Al, K, Ca, Mn, Fe, Ni, Cu, Ga, Se, Sr, Ba, Rb and Pb**<br>mg L<sup>-1</sup> | **% grass covered shore**<br><10, 11-20, 21-30, 31-40, 41-50, 51-60, 61-70, 71-80, 81-90, 91-100<br>**% grass covered slopes**<br><10, 11-20, 21-30, 31-40, 41-50, 51-60, 61-70, 71-80, 81-90, 91-100 |  |

**List S1.**
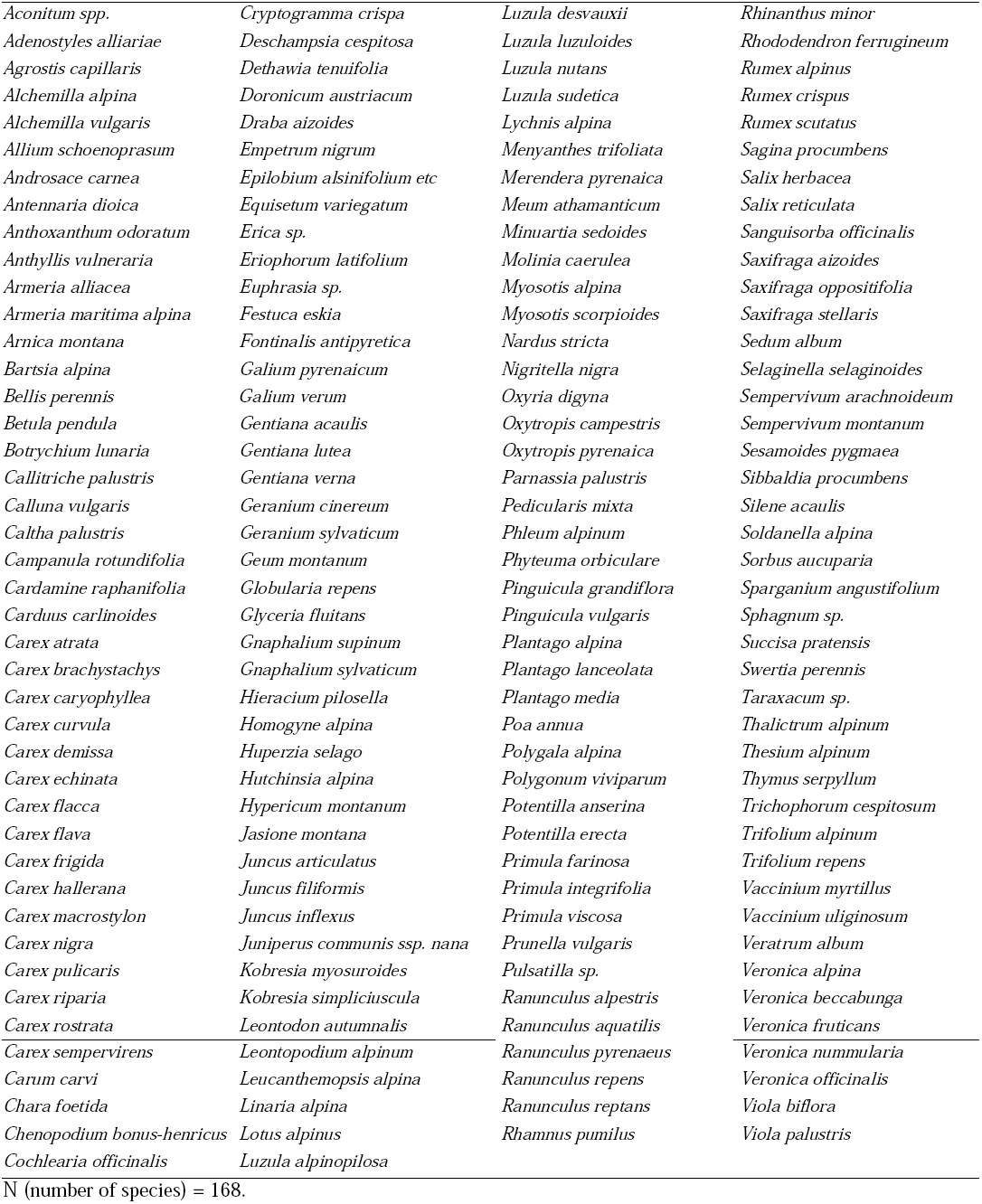
The taxonomical composition of riparian vegetation in the 189 central Pyrenean lakes and ponds surveyed for this study.

## REFERENCES

Alam SM. 1999. Nutrient uptake by plants under stress conditions. In: Pessarakli M. (Ed), Handbook of plant and crop stress, 2^nd^ ed. revised and expanded. Marcel Dekker, New York, USA, pp. 285–314.

Austin MP, Smith TM. 1989. A new model for the continuum concept. Vegetatio 83: 35–47.

Báez JC, Real R, Mario-Vargas J, Flores-Moya A. 2005. Chorotypes of seaweeds from the western Mediterranean Sea and the Adriatic Sea: An analysis based on the genera Audouinella (Rhodophyta), Cystoseira (Phaeophyceae) and Cladophora (Chlorophyta). Phycol Res 53: 255–265.

Baker WL. 1989. Macro- and micro-scale influences on riparian vegetation in Western Colorado. Ann Assoc Am Geogr 79(1): 65– 78.

Bastian M, Heymann S, Jacomy M. 2009. Gephi: an open source software for exploring and manipulating networks. International AAAI Conference on Weblogs and Social Media.

Baroni-Urbani C, Ruffo S, Vigna-Taglianti A. 1978. Materiali per una biogegrafia italiana fondata su alcuni generi di coleotteri cicindelidi, carabidi e crisomelidi. Estr Mem Soc Ent Ital 56: 35–92.

Beeson KC. 1959. Magnesium in soil-sources, availability and zonal distribution. In: Magnesium and agriculture, Proceedings of the symposium; Anderson GC, Jencks EM, Horvath DJ (Eds), West Virginia University, Morgantown: 1–11.

Boyce R. 2008. Fuzzy set ordination web page. Northern Kentucky University. Accessed: August 2009 from http://www.nku.edu/~boycer/fso/.

Boyce RL, Ellison PC. 2001. Choosing the best similarity index when performing fuzzy set ordination on binary data. J Veg Sci 12:711–20.

Buma B, Hennon PE, Harrington CA, Popkin JR, Krapek J, Lamb MS, Oakes LE, Saunders S, Zeglen S. 2016. Emerging climate-driven disturbance processes: widespread mortality associated with snow-to-rain transitions across 10° of latitude and half the range of a climate-threatened conifer. Global Change Biology. DOI: 10.1111/gcb.13555

Castillo-Jurado M. 1992. Morfometria de los lagos. Una aplicacion a los lagos del Pirineo. PhD thesis, University of Barcelona (in Spanish).

Catalan J, Camarero L, Felip M, Pla S, Ventura M, Buchaca T, Bartumeus F, De Mendoza G, Miró A, Casamayor EO, Medina-Sánchez JM, Bacardit M, Altuna M, Bartrons M, De Quijano DD. 2006. High mountain lakes: extreme habitats and witnesses of environmental changes. Limnetica 25(1-2): 551–584.

Couto W, Sanzonowics C, Leite GG. 1983. Effect of excess water in an oxisol on ammonium, nitrate, iron and manganese availability and nutrient uptake of two tropical forage species. Plant Soil 73: 159–166.

Dufrene M, Legendre P. 1997. Species assemblages and indicator species: the need for a flexible asymmetrical approach. Ecol Monogr 67(3):345–366.

EEA (European Environment Agency). 2001. Biogeographical regions, Europe 2001. Available online at http://www.eea.europa.eu/data-and-maps/figures/biogeographical-regions-europe-2001. Accessed February 04, 2010.

Engelhardt BM, Chambers JC, Weisberg PJ. 2015. Geomorphic predictors of riparian vegetation in small mountain watersheds. Journal of Plant Ecology 8 (6), 593–604.

Fattorini, S., 2015. On the concept of chorotype. Journal of Biogeography, 42, 2246–2251.

Fitter RSR, Fitter A, Farrer A. 1984. Grasses, sedges, rushes and ferns of Britain and Northern Europe. Collins, London, UK.

García-Rollán M. 1985. Claves de la flora de España (peninsula y Baleares). Vol.II: dicotiledoneas (l-z) / monocotiledoneas. Mundi-Prensa [in Spanish].

Gómez D, Sesé JA, Villar L. 2003. The vegetation of the alpine zone in the Pyrenees. In: Alpine biodiversity in Europe, Nagy L, Grabherr G, Körner Ch, Thompson DBA (Eds). Ecol Studies 167: 85–92.

Good BJ, Patrick Jr, WH. 1987. Gas composition and respiration of water oak (Quercus nigra L.) and green ash (Fraxinus pennsylvanica Marsh.) roots after prolonged flooding. Plant Soil 97: 419–27.

Gregory SV, Swanson FJ, McKee WA, Cummins KW. 1991.An ecosystem perspective of riparian zones: focus on links between land and water. BioScience 41(8): 540–549.

Grey-Wilson C, Blamey M. 1979. The alpine flowers of Britain and Europe. Collins, London, UK.

Grey-Wilson C, Blamey M. 1995. The alpine flowers of Britain and Europe. Collins, London, UK.

Gruber M. 1992. Schéma des séries dynamiques de végéta%on des Hautes-Pyrénées (Pyrénées centrales francaises). Bot Complutensis 17: 7–21.

Hagmann P. 2005. From Diffusion MRI to Brain Connectomics. Science 3230:127.

Hengeveld R. 1990. Dynamic biogeography. Cambridge University Press, Cambridge.

Hennig C. 2005. A method for visual cluster validation. In: Weihs C, Gaul W (eds). Classification - The Ubiquitous Challenge. Springer, Heidelberg 2005, 153–160.

Kabata-Pendias A, Pendias H. 2001. Trace elements in soils and plants. 3^rd^ ed. CRC Press LLC, USA.

Kaufman L, Rousseeuw PJ. 1990. Finding groups in data: an introduction to cluster analysis. Wiley, New York.

Keller F, Goyette S, Beniston M. 2005. Sensitivity analysis of snowcover to climate change scenarios and their impact on plant habitats in alpine terrain. Clim Change 72: 299–319.

Kernan M, Ventura M, Bitušík P, Brancelj A, Clarke G, Velle G, Raddum GG, Stuchlík E, Catalan J. 2009. Regionalisation of remote European mountain lake ecosystems according to their biota: environmental versus geographical patterns. Freshwat Biol 54 (12): 2470–93.

Khamis K, Hannah DM, Brown LE, Tiberti R, Milner a. M. 2014. The use of invertebrates as indicators of environmental change in alpine rivers and lakes. Sci Total Environ 493:1242– 54.

Körner Ch. 1999. Alpine plant life: functional plant ecology of high mountain ecosystems. Springer, Berlin Heidelberg, p. 343.

Kovalchik BL, Chitwood LA. 1990. Use of geomorphology in the classification of riparian plant associations in mountainous landscapes of central Oregon, U.S.A. For Ecol Manage 33/34: 405–418.

Kreyling J, Jentsch A, Beier C. 2014. Beyond realism in climate change experiments: gradient approaches identify thresholds and tipping points. Ecol Lett 17: 125–e1.

Larsen, I.J., Montgomery, D.R. & Greenberg, H.M., 2014. The contribution of mountains to global denudation. Geology 42(6), pp.527–530.

Lewin J, Macklin MG. 1987. Metal mining and floodplain sedimentation in Britain. In: Gardiner V (ed). International geomorphology, part 1. Chichester, Wiley, p. 1009–27.

Loucks OL. 1962. Ordinating forest communities by means of environmental scalars and phytosociological indices. Ecol Monogr 32: 137–166.

Merritt DM, Scott ML, Poff NL, Auble GT, Lytle DA. 2010. Theory, methods and tools for determining environmental flows for riparian vegetation: riparian vegetation-flow response guilds. Freshwat Biol 55: 206–225.

Minot JM, Carrillo E, Font X, Carreras J, Ferré A, Masalles RM, Soriano I, Vigo J. 2007. Altitude zonation in the Pyrenees. A geobotanic interpretation. Phytocoenologia 37 (3–4): 371–398.

Oksanen J, Kindt R, Legendre P, O’Hara B, Simpson GL, Solymos P, Stevens MHH, Wagner H. 2009. Vegan: community ecology package. R package version 1.15-4. http://CRAN.Rproject.org/package=vegan.

Parker BR, Vinebrooke RD, Schindler DW. 2008. Recent climate extremes alter alpine lake ecosystems. Proc Natl Acad Sci U S A 105:12927–31.

Price MF (ed.). 2004. Conservation and sustainable development in mountain areas. IUCN, Gland, Switzerland and Cambridge, UK. 29 pp.

R Core Development Team. 2005. R: a language and environment for statistical computing. R Foundation for Statistical Computing, Vienna, Austria (http://www.R-project.orgi).

Roberts DW. 1986. Ordination on the basis of fuzzy set theory. Vegetatio 66: 123–131.

Roberts DW. 2007(a). FSO: fuzzy set ordination. R package version 1.0-1 (http://cran.Rproject.orgi).

Roberts DW. 2007(b). LabDSV: ordination and multivariate analysis for ecology. R package version 1.3-0 (http://cran.R-project.orgi).

Roberts DW. 2008. Statistical analysis of multidimensional fuzzy set ordinations. Ecology 89:1246–1260.

Roberts DW. 2009(a). Comparison of multidimensional fuzzy set ordination with CCA and DBRDA. Ecology 90(9), 2622–34.

Roberts DW. 2009(b). Fuzzy set ordination, R labs for vegetation ecologists, Montana State University. http://ecology.msu.montana.edu/labdsv/R/labs/lab11/lab11.html. Accessed August 2009.

Sporns O. 2006. Small-world connectivity, motif composition, and complexity of fractal neuronal connections. BioSystems 85:55–64.

Storkey J, Macdonald AJ, Poulton PR, Scott T, Köhler IH, Schnyder H, Goulding KWT, Crawley MJ. 2015. Grassland biodiversity bounces back from long-term nitrogen addition. Nature, 528(7582), 401–404.

Thioulouse J, Chessel D, Doledee S, Olivier JM. 1997. ADE-4 A multivariate analysis and graphical display software. Stat Comput 7: 75–83.

Thompson R, Kamenik C, Schmidt R. 2005. Ultra-sensitive Alpine lakes and climate change. J Limnol 64:139–52.

Verardo DJ, Froelich PN, McIntyre A. 1990. Determination of organic carbon and nitrogen in marine sediments using the Carlo Erba NA-1500 Analyzer. Deep-Sea Res. Part A, 37: 157–165.

Whittaker RH. 1967. Gradient analysis of vegetation. Biol Rev 42: 207–264.

Williamson C, Saros JE, Vincent WF, Smol JP. 2009. Lakes and reservoirs as sentinels, integrators, and regulators of climate change. Limnol Oceanogr 54: 2273–82.

Wong CC, Wu SC, Kuek C, Khan AG, Wong MH. 2007. The role of mycorrhizae associated with vetiver grown in Pb-/Zn-contaminated soils: Greenhouse study. Restor Ecol 15(1): 60–67.

Zaharescu DG, Hooda PS, Fernandez J, Soler AP, Burghelea CI. 2009. On the arsenic-source mobilisation and its natural enrichment in the sediments of a high mountain cirque in the Pyrenees. J Environ Monit 11: 1973–1981.

Zaharescu DG, Burghelea CI, Hooda PS, Polyakov V, Palanca-Soler A. 2016(a). Climate change enhances the mobilisation of naturally occurring metals in high altitude environments. The Science of the Total Environment 560-561: 73–81.

Zaharescu DG, Hooda PS, Burghelea CI, Palanca-Soler A. 2016(b). A multiscale framework for deconstructing the ecosystem physical template of high altitudes lakes. Ecosystems 19 (6): 1064–1079.

Zaharescu DG, Burghelea CI, Hooda PS, Lester RN, Palanca-Soler A. 2016(c). Small lakes in big landscape: Multi-scale drivers of littoral ecosystem in alpine lakes. The Science of the Total Environment 551–552: 496–505.

